# Right-lateralized prefrontal signature with successful Stroop-like interference control in early childhood

**DOI:** 10.64898/2026.07.16.738912

**Authors:** Ryuta Kuwamizu, Nozomi Yamamoto, Kota Otani, Yusuke Moriguchi

**Affiliations:** Institute of Health and Sport Sciences, University of Tsukuba, Ibaraki, Japan; Graduate School of Letters, Kyoto University, Kyoto, Japan; Graduate School of Human Sciences, The University of Osaka, Osaka, Japan

## Abstract

The Stroop effect is a canonical measure of executive control, yet its neural architecture has been defined largely in literate children and adults, where left frontal mechanisms have long been implicated in resolving verbal conflict. How the developing brain supports Stroop-like interference control before literacy and stable verbal responding are established remains unknown. Here we show that successful interference control in early childhood is associated with the selective engagement of right lateral prefrontal regions. We used multichannel functional near-infrared spectroscopy to measure prefrontal hemodynamics in 94 children aged 35–79 months during a color-pointing Stroop-like task, in which children pointed to colors in response to spoken color names. Stroop-like conflict elicited broad activation across bilateral lateral prefrontal regions, a conflict response already present from around 3 years of age and did not show a detectable age-related increase. By contrast, individual differences in accuracy under conflict were selectively associated with greater activation in the right dorsolateral and right rostrolateral prefrontal cortices, independent of age. These findings suggest that right lateral prefrontal regions play an important role in successful interference control in early childhood, indicating that preschool Stroop-like control is not merely a weaker form of the adult left-lateralized system.

## Introduction

Executive function and self-regulation in preschool predict later academic achievement, social adjustment, and mental and physical health^1–3^. Inhibitory control, a core component of executive function, is the ability to suppress an automatic response when it conflicts with current goals^4,5^, which improves increasingly rapidly across the preschool years^6^. The question of how the developing brain supports this rapid gain, however, remains unresolved. When a preschool child succeeds in overriding a prepotent response, is that success already carried by a functionally organized prefrontal system, and if so, is that system organized as in the adult brain or along different lines?

The Stroop task is a classic and widely used measure of interference control, indexing the conflict that arises when language and color information compete^7,8^. Resolving Stroop conflict draws on a distributed frontocingulate–parietal network in which the anterior cingulate cortex signals conflict and, in concert with the parietal cortex, the lateral prefrontal cortex (LPFC) implements control^9–13^. Within this network, the left LPFC has consistently been the most strongly recruited region and most closely tied to performance^10,12,14^. An early study on brain lesions showed that damage to the left frontal lobe produces a growing deficit as Stroop interference increases, most markedly when verbal factors are involved^15^, shaping the lasting view of Stroop interference control as a predominantly left-hemispheric, verbally mediated function. This left-weighting persists, remaining consistent across both functional magnetic resonance imaging (fMRI; including coordinate-based meta-analyses and recent studies^10,12,16^) and functional near-infrared spectroscopy (fNIRS)^17–19^ (but some studies have also observed bilateral engagement and noted a functional contribution of the right hemisphere^16,20^) studies. More specifically, fNIRS studies using interhemispheric difference measures have linked relatively greater left-than-right DLPFC activation to less Stroop interference in both young and older adults^21,22^. Indeed, improvements or decrements in Stroop task performance induced by physical exercise, music, and a hypoxia environment are likewise accompanied by corresponding changes in left lateral prefrontal activation^23–27^. The importance of the left LPFC for interference control is thus well established in the adult brain. Developmental neuroimaging studies have shown that, from around the age of seven onward, activity in a frontoparietal network including the left LPFC increases with age during Stroop interference^14,28^, suggesting that this left-lateralized functional organization is progressively strengthened over the school years.

This leaves open what happens earlier, in preschool-aged children, whose language is still immature and who cannot yet perform the classic verbal Stroop task. Whether this left-dominant organization also applies to the developing brain is unknown. Outside the Stroop task, the developmental literature more consistently points to earlier functional maturation of the right hemisphere in infancy and early childhood. In infancy, the right cortex appears to develop ahead of the left leading to right-lateralized task-related and resting state activation^29^. The same right-hemisphere bias appears in preschoolers with non-Stroop executive function tasks such as the Dimensional Change Card Sort (DCCS), where successful performance predominantly recruits the right LPFC^30–34^, and right-lateralization strengthens as performance improves with age^30,31^.

Whether early Stroop interference control depends on the left-dominant, verbally mediated system of the mature Stroop task or on this right-lateralization specific to early development remains an open question. The classic color-word Stroop task demands literacy and stable verbal responding and, thus, is not straightforward in its administration to preschool children in its original form^5,35^. Therefore, developmental work relies on age-appropriate variants^36^. Despite superficial differences from the classic color-word Stroop, these variants are designed to preserve its cognitive core, namely prioritizing one verbal response over competing alternatives while suppressing the more prepotent one; they remain, however, Stroop-like rather than the Stroop task itself, a distinction we develop further in the Discussion. These variants produce reliable interference in participants from the age of two to three years^34,36–40^, but the neural basis remains unclear, with prefrontal activation reported inconsistently (i.e., activation or deactivation, right or left). Moreover, the relationship between interference and successful task performance is still undetermined^34,37–40^

The present study addressed this issue using multichannel fNIRS with 94 children aged 35– 79 months during a color-pointing Stroop-like task that yields robust behavioral interference^40^. Our group has previously shown that prefrontal activation in early childhood tracks performance in cognitive shifting tasks^30–32^. We asked whether conflict elicits lateral prefrontal activation, whether this activation is associated with successful interference control independently of age, and whether any such brain–behavior association is left-dominant, as in adults, or right-lateralized. Here we show that the Stroop-like conflict recruited the bilateral LPFC from the youngest age tested, yet only right dorsolateral and right rostrolateral activation predicted successful performance under conflict, independently of age. These results indicate that preschool interference control is supported not by broader recruitment but by a selective, right-lateralized functional specialization within an already engaged LPFC.

## Results

### Behavioral performance

First, to establish that our Stroop-like task elicited conflict-based interference, we compared accuracy between conditions. Accuracy was significantly lower for incongruent than congruent trials (0.750 ± 0.260 vs. 0.954 ± 0.053; Wilcoxon *W* = 4,131, *P* < 0.001) (Fig. 1). Accuracy improved with age in both the incongruent (*r* = 0.540, *P* < 0.001) and congruent (*r*= 0.347, *P* < 0.001) conditions, and the interference score (incongruent accuracy minus congruent accuracy) likewise decreased with age (*r* = 0.506, *P* < 0.001), consistent with the protracted development of inhibitory control at this age. Together, these indices confirm that the task engaged conflict processing in participants 3 to 6 years of age.

**Figure 1.**
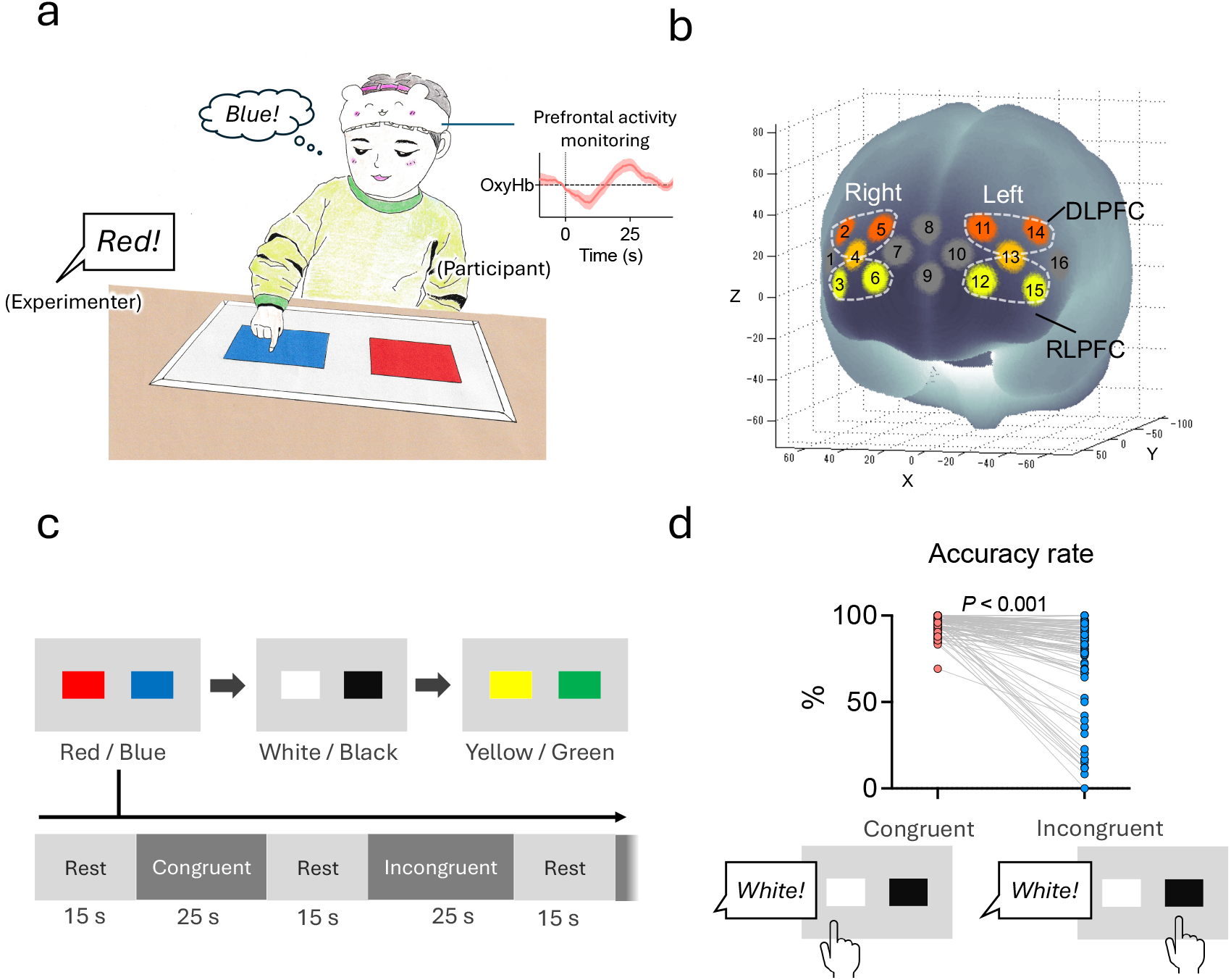
Experimental design, behavioral results, and fNIRS channel layout for the color-pointing Stroop-like task. (A) Example of the age-appropriate color-pointing Stroop-like variant. Children listened to spoken color labels and responded by pointing, without reading words or producing verbal color names, while prefrontal activity was monitored using fNIRS. In incongruent trials, children selected the rule-relevant color response while suppressing the competing linguistically cued named-color response, preserving the Stroop-like requirement to prioritize one color-related response over another under conflict. (B) Spatial layout of the 16-channel fNIRS probe array registered to MNI standard space. Channels are numbered 1–16; highlighted channels indicate those assigned to each region of interest. (C) Three color pairs (red/blue, white/black, yellow/green) were presented sequentially on A3-sized laminated cards. Each condition comprised a 15-s rest phase followed by a 25-s task phase. (D) Individual accuracy rates are shown for congruent (red) and incongruent (blue) trials, with lines connecting data from the same child. Accuracy was significantly lower in the incongruent than congruent condition (P < 0.001, Wilcoxon signed-rank test).

### Bilateral PFC recruited by conflict from 3 years of age

We next examined whether this behavioral interference was accompanied by increased prefrontal recruitment, and whether such recruitment was already present across 3-to 6-year-olds (Fig. 2). A linear mixed-effects model on oxy Hb change (Condition × Hemisphere × Region, with random intercepts for participant) revealed a robust main effect of condition (*F*(1, 640.5) = 74.74, *P* < 0.001), with all four prefrontal regions of interest (ROIs) showing stronger activation during incongruent than congruent trials (left rostrolateral (RL) PFC: *t*(641.0) = 4.59; right RLPFC: *t* = 3.30; left dorsolateral (DL) PFC: *t* = 4.09; right DLPFC: *t* = 5.32; all *P* ≤ 0.001). Signals were overall larger in the right than left hemisphere (*F*(1, 641.9)= 5.02, *P* = 0.025) and in dorsal than in rostral regions (*F*(1, 641.9) = 26.64, *P* < 0.001), but no interaction involving condition reached significance (all *P* > 0.22), indicating a conflict response in the bilateral PFC. Importantly, adding age to the model did not reveal developmental modulation (main effect: *F*(1, 91.1) = 0.38, *P* = 0.538; age × condition: *F*(1, 639.5) = 1.98, *P* = 0.160), nor did sex (*F*(1, 90.5) = 0.84, *P* = 0.363). Model-derived estimates confirmed that the conflict-related increase was already significant at 36 months (estimate =−0.012, *t*(640) = −2.55, *P* = 0.011) and remained significant at 48 months (*t* = −5.27, *P* < 0.001), 60 months (*t* = −8.76, *P* < 0.001), and 72 months (*t* = −6.63, *P* < 0.001). The prefrontal conflict response is therefore present from 3 years of age and does not measurably increase in magnitude across the preschool years.

**Figure 2.**
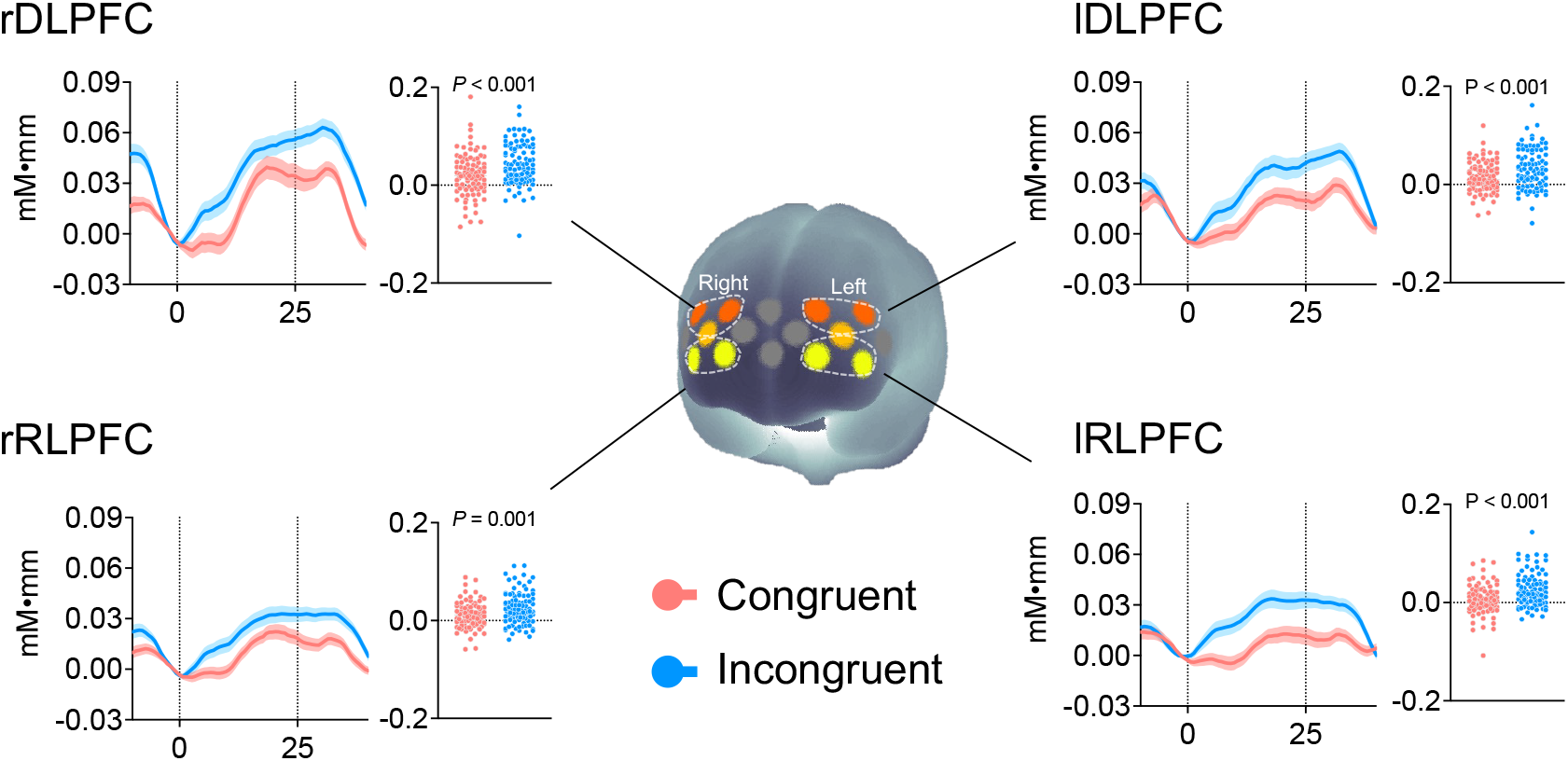
Bilateral lateral prefrontal cortex (LPFC) is recruited by conflict. Mean oxy-Hb change (mM·mm) during congruent and incongruent trials in each prefrontal region of interest (left and right dorsolateral PFC (DLPFC), left and right rostrolateral PFC (RLPFC)). Error bars denote SEM. n = 94 children aged 35–79 months.

### Right LPFC selectively tracks individual differences in conflict performance

To test whether prefrontal recruitment was functionally coupled to behavior in a manner specific to conflict processing, we examined brain–behavior correlations separately for each condition (Fig. 3). Incongruent activation correlated positively with incongruent accuracy in the right DLPFC (*r* = 0.359, *P* < 0.001, *P*_FDR = 0.001) and the right RLPFC (*r* = 0.343, *P* < 0.001, *P*_FDR = 0.001), whereas left-hemisphere correlations were weaker, with the left DLPFC showing only a marginal effect (*r* = 0.221, *P* = 0.038, *P*_FDR = 0.050) and the left RLPFC failing to reach significance (*r* = 0.091, *P* = 0.381). These right-hemisphere associations under conflict were robust to demographic and behavioral confounds: the original correlations remained after partialling out age (r DLPFC: *r* = 0.294, *P* = 0.004, *P*_FDR = 0.008; r RLPFC: *r* = 0.352, *P* < 0.001, *P*_FDR = 0.002), and were essentially unchanged after additionally partialling out sex (rDLPFC: *r* = 0.300, *P* = 0.004, *P*_FDR = 0.007; r RLPFC: *r* = 0.356, *P* < 0.001, *P*_FDR = 0.002) or the number of completed incongruent trials (r DLPFC: *r* = 0.287, *P* = 0.006, *P*_FDR = 0.011; r RLPFC: *r* = 0.357, *P* < 0.001, *P*_FDR = 0.002). These associations were also unchanged after controlling for hair color and hand use (Supplementary Note 1). By contrast, congruent activation was unrelated to congruent accuracy in every ROI (all |*r*| < 0.06, all *P* > 0.57). The interference index, computed as incongruent minus congruent for both accuracy and oxy-Hb, also yielded positive associations in the same right-hemisphere ROIs (r RLPFC: *r* = 0.257, *P* = 0.012, *P*_FDR = 0.049; r DLPFC: *r* = 0.225, *P* = 0.029, *P*_FDR = 0.058).

**Figure 3.**
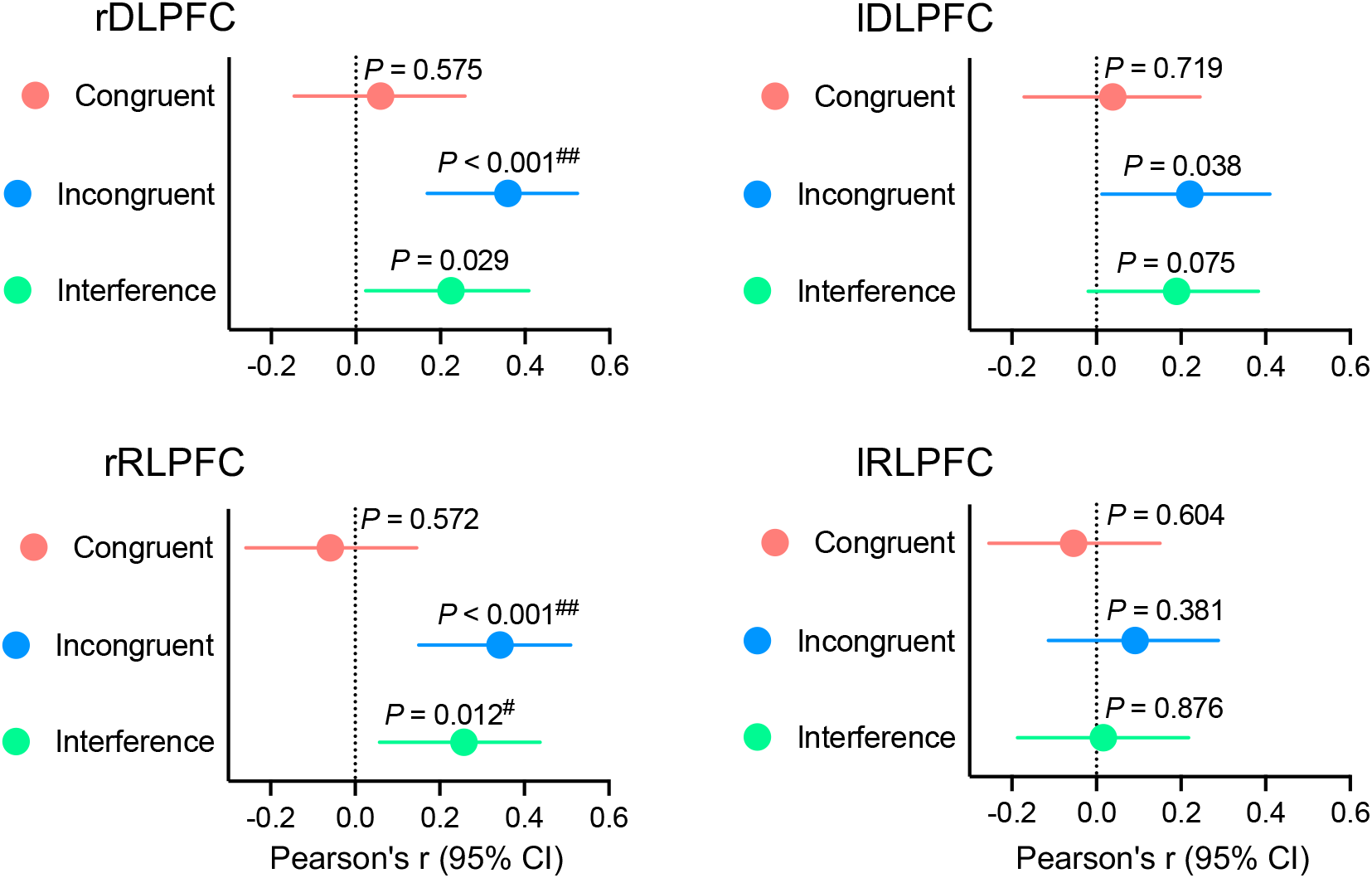
Right lateral prefrontal activation selectively predicts conflict performance. Forest plots showing Pearson correlations between prefrontal oxy-Hb responses and behavioral accuracy across four ROIs (right (r) and left (l) dorsolateral prefrontal cortex (DLPFC) and right and left rostrolateral prefrontal cortex (RLPFC)) and three conditions: Incongruent (oxy-Hb vs. accuracy in the incongruent condition), Congruent (oxy-Hb vs. accuracy in the congruent condition), and Interference (incongruent-minus-congruent oxy-Hb difference vs. accuracy difference). Dots indicate Pearson’s r; horizontal bars indicate 95% confidence intervals (CI). Dashed lines indicate r = 0. ^#^P_FDR < 0.05, ^##^P_FDR < 0.01 (FDR-corrected across the four ROIs within each condition). The brain–behavior coupling under conflict was selective to the right-hemisphere prefrontal cortex and absent in the congruent condition, indicating condition- and hemisphere-specific neural recruitment for successful interference control. n = 94 children aged 35–79 months.

As exploratory analyses, because both significant effects were located in the right hemisphere, we next asked whether this lateralization was itself reliable (Fig. 4). A hemispheric asymmetry index (right minus left oxy-Hb)^21^ computed for the RLPFC was positively associated with incongruent accuracy (*r* = 0.309, *P* = 0.002, *P*_FDR = 0.005), indicating that children who more strongly engage the right relative to the left RLPFC during conflict tend to perform more accurately; the same RLPFC asymmetry index, computed on the interference contrast, was likewise associated with the interference accuracy score (*r* = 0.300, *P* = 0.003, *P*_FDR = 0.007). The DLPFC asymmetry index, by contrast, showed a relationship in the same direction for incongruent trials but did not reach significance (*r =* 0.179, *P* = 0.093) and was absent for the interference contrast (*r* = 0.036, *P* = 0.739). Together, the condition contrast and the hemispheric pattern indicate that conflict-related individual differences in performance are tracked specifically by right-hemisphere prefrontal recruitment, with the RLPFC asymmetry providing the strongest lateralization signal.

**Figure 4.**
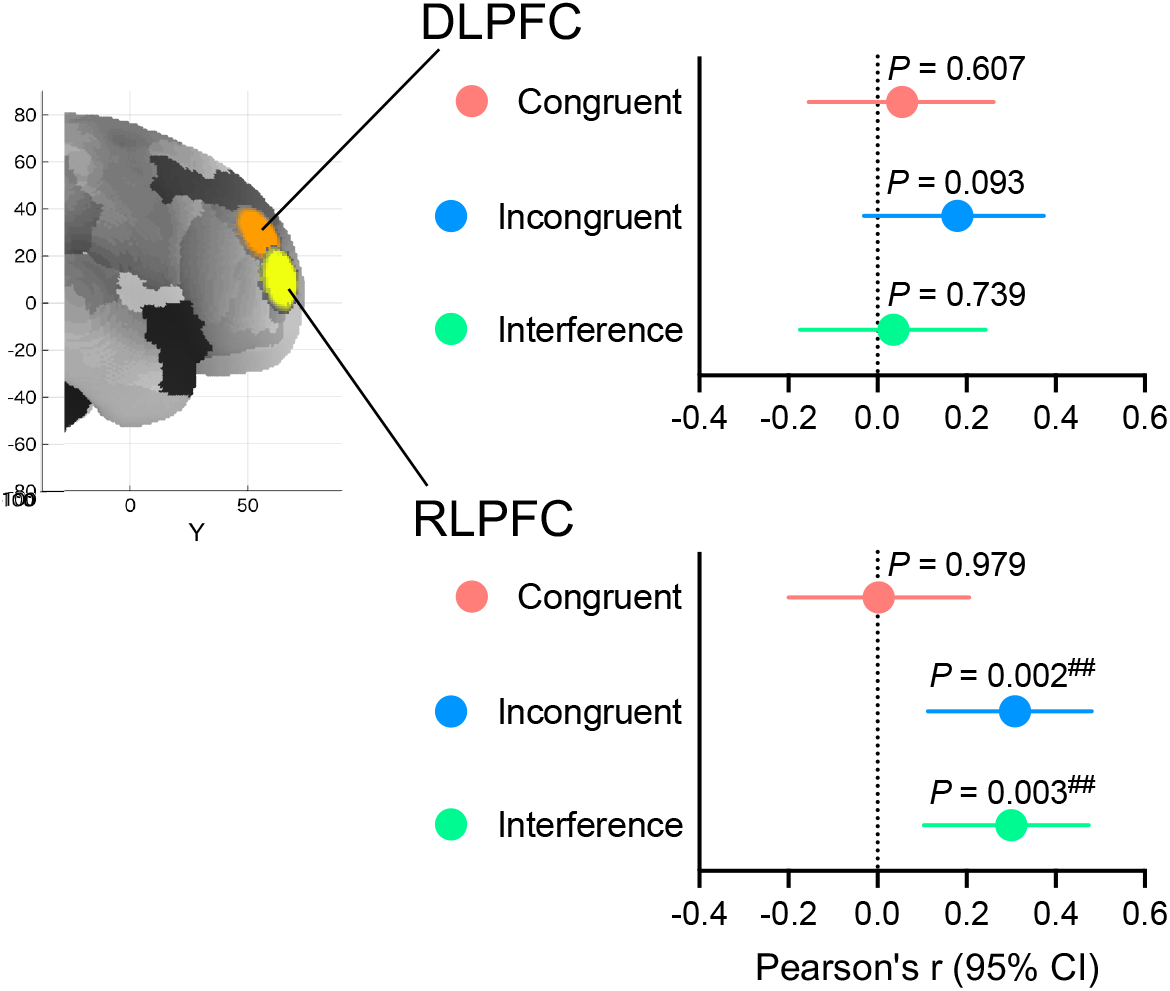
Hemispheric lateralization of prefrontal activation and conflict performance. Forest plot showing Pearson correlations between the prefrontal lateralization index (right-minus-left hemispheres oxy-Hb) and behavioral accuracy, computed separately for dorsolateral prefrontal cortex (DLPFC) and rostrolateral prefrontal cortex (RLPFC) across three conditions (Incongruent, Congruent, and Interference; defined as in Fig. 3). Dots indicate Pearson’s r; horizontal bars indicate 95% confidence intervals (CI). The dashed lines indicate r = 0. ^#^P_FDR < 0.05, ^##^P_FDR < 0.01 (FDR-corrected across the two ROIs within each condition).

## Discussion

Preschool-age Stroop-like interference control is not simply an immature form of the adult left-lateralized Stroop system. In 94 children aged 35 to 79 months, Stroop-like conflict elicited broad bilateral lateral prefrontal activation that was already present from the youngest age tested and showed no detectable age-related strengthening across the preschool years. Yet individual differences in successful conflict performance were selectively associated with right DLPFC and right RLPFC activation during incongruent trials. These findings reveal an early prefrontal architecture in which broad conflict-related recruitment coexists with a right-lateralized performance signal before children can perform the classic literacy-based color– word Stroop task.

This pattern reframes a contested developmental timeline. Earlier neuroimaging work suggested that a behaviorally meaningful prefrontal contribution to Stroop interference becomes reliably detectable only around age 7 and strengthens gradually into adolescence^14,28^. Subsequent fNIRS studies of child-friendly Stroop variants reported preschool DLPFC activation in some cases^34,37^ but not others^39^, leaving the literature mixed rather than settled. Although these variants are not the classic color-word Stroop task itself, they preserve a related Stroop-like control demand: selecting a rule-relevant color response while suppressing a competing response cued by verbal-semantic color information. Our data speak to this debate by indicating that an organized prefrontal contribution to interference control is present earlier than the late-maturation account assumed^14,28^. Behavioral improvement across the preschool years therefore likely reflects gains in the efficiency and stability with which an already organized circuit is engaged, rather than the gradual emergence of a new prefrontal contribution.

The right-lateralized character of the performance signal, present against a backdrop of bilateral conflict-evoked activation, is particularly noteworthy. This interpretation does not treat the present color-pointing task as the classic color-word Stroop task itself; rather, it treats it as a Stroop-like variant that preserves a related control demand. In adults, the classic color-word Stroop task typically recruits left lateral prefrontal regions, reflecting the need to resolve conflict between competing color-related verbal-semantic representations^10–12^. In the present task, conflict arose between the prepotent response to the named color and the rule-relevant response to the opposite color, requiring children to prioritize one verbally cued color-response representation while suppressing another. Importantly, the typical left-prefrontal pattern has also been reported in Stroop variants using manual responses, indicating that it cannot be attributed solely to overt speech production or response mode in adults^10,41^. Recently, Okayasu et al.^10^ reported that this left lateralization of the lateral prefrontal cortex emerges under both manual (button-press) and verbal responses, but not during non-verbal conflict tasks such as the flanker task, indicating that it reflects the verbal-semantic nature of the conflict rather than the response mode or the mere presence of conflict. Against this background, the right-lateralized brain–behavior association observed here is unlikely to be explained simply by the pointing response and may instead reflect a developmentally early form of control for resolving verbally cued color-response conflict.

This developmental interpretation also converges with prior developmental fNIRS studies of other executive-function paradigms, such as rule-switching, which have repeatedly localized preschool and infant brain–behavior coupling to the right LPFC^29–33^. This convergence across surface-different tasks that share the need to prioritize a goal-relevant response while suppressing a prepotent alternative suggests that the right-hemisphere bias in PFC activation seen here may not be paradigm-specific but a broader property of this developmental period. This pattern may reflect a developmental sequence in which the right LPFC comes online first. The right hemisphere has also been reported to mature earlier than the left functionally^42^. Consistent with this, young children who master rule-switching earliest engage the right LPFC^30^, whereas those who succeed about a year later engage the left^43^. We speculate that the LPFC is engaged from early in development, but that its two hemispheres follow different developmental timetables: the right LPFC may become functionally important first, consistent with the right-lateralized brain–behavior association observed in the present study, while the left hemisphere develops more gradually thereafter and comes to support performance as well. From this view, the left-lateralized, verbally mediated organization characteristic of the adult Stroop task emerges progressively, as left frontoparietal engagement during Stroop interference increases with age^14,28,44^, yielding the more bilateral, task-tuned system of the mature brain. Because several cortical measures are already left–right asymmetric by one to six years of age^45^, this early lateralization is perhaps less the appearance of asymmetry from nothing than a shifting left–right balance in task engagement, strategy, and network participation, built upon an already-asymmetric structural foundation.

Examining the regions in more detail, the RLPFC showed clearer right-dominance, with its asymmetry index reliably predicting performance (Fig. 4). This dissociation is notable because interference-related prefrontal activation in adults is more commonly localized to the dorsolateral region of the PFC^10,12^, whereas here it was the rostrolateral region that carried the more reliable lateralized brain–behavior association. The RLPFC is known to be involved in higher-order cognitive functions such as relational reasoning and abstract thinking^46–49^, rather than the more basic conflict resolution usually associated with the DLPFC. In the current results, the stronger lateralization signal in the RLPFC may reflect the possibility that the present Stroop-like task was particularly higher-order demanding for preschool children: young children were required to maintain the task rule, suppress a prepotent stimulus-driven response, and select the appropriate response based on an abstract mapping. These demands align with the proposed roles of the RLPFC in higher-order control^46–49^, suggesting that this region can be recruited early in development when task demands require high-level control.

More broadly, our findings pertain to the long-standing question of how the developing PFC supports cognitive control: does development bring stronger overall recruitment, or a shift from diffuse to more focal, specialized engagement^50,51^? In our data, even younger and lower-performing children already showed robust prefrontal activation under conflict; what distinguished successful performance was not greater overall recruitment but selectively higher activation in the right DLPFC and right RLPFC. The developmental signature of successful interference control is therefore functional specialization within an already engaged system, identifying a candidate neural locus through which early experience and intervention may shape lifelong self-regulation: not by recruiting the PFC more broadly, but by sharpening the functional specialization that supports successful control. That said, because the present task is Stroop-like rather than the classic color-word Stroop task and our sample spanned only 3 to 6 years of age, this interpretation should be tested across a broader age range before it is generalized.

These findings carry methodological implications that reach beyond preschool conflict processing itself. The combination of a Stroop task with fNIRS has become a widely used approach for assessing how acute and chronic interventions, such as physical exercise, music training, mindfulness, and educational programs, enhance executive function in other populations, because it yields a behaviorally validated and minimally invasive readout of prefrontal executive function^23,25,27,52–54^. Yet this paradigm has remained largely out of reach in the very developmental window in which inhibitory control gains most rapidly and in which early interventions carry the largest leverage on later academic, health, and social outcomes^1,3^. Critically, the present paradigm proved highly feasible even at this young age range: more than 99% of the measured trials and channels were retained for analysis, demonstrating that high-quality fNIRS data can be obtained from 3-to 6-year-olds. By demonstrating that a child-friendly color-pointing Stroop variant produces a robust, conflict-specific, right-lateralized brain–behavior signal in 3-to 6-year-olds, the present study extends this widely used approach into early childhood.

Several limitations should be noted. First, although the task was designed to tax interference control, contributions from working memory and rule maintenance cannot be fully ruled out. Moreover, the present color-pointing task is not identical to the classic color-word Stroop task as children listened to spoken color labels and pointed to colors without reading words or producing verbal color names. Thus, verbal processing was still involved, but differences in stimulus format and response demands may have influenced the balance of cognitive processes engaged by the task^10,20^. We therefore regard our paradigm as Stroop-like rather than as a direct instantiation of the classic color-word Stroop task. Because the sample was restricted to 3-to 6-year-olds, the present data cannot directly determine how this early right-lateralized organization relates to the adult left-lateralized Stroop system. Future studies should extend this color-pointing paradigm to older children and adults and, ideally, administer the classic color-word Stroop task and the Stroop-like variant to the same individuals. The color-pointing format nonetheless avoids the arbitrary label–stimulus mappings of the Day–Night task and provides a more direct measure of inhibitory control than some alternatives^55^. Second, we did not measure trial-level reaction times, because manual pointing in young children does not yield the necessary temporal precision; we therefore relied on accuracy as the primary index of conflict cost and confirmed that the brain–behavior associations were essentially unchanged after controlling for the number of completed trials within each block (an approximate index of speed). Finally, fNIRS interpretation can in principle be affected by extracerebral and systemic physiological contributions^56^; we mitigated this by restricting recordings to the frontal scalp, where hair-related signal loss is minimal, and by applying hemodynamic modality separation to suppress residual non-neural components^57^.

In conclusion, Stroop-like conflict already engages the bilateral LPFC in early childhood, yet successful interference control is marked by a right-lateralized prefrontal performance signal. These findings suggest that interference control in young children is supported by right-sided selectivity within an already engaged prefrontal architecture, rather than by broader frontal recruitment alone. This pattern raises the possibility that early Stroop-like control is not merely a weaker precursor of the adult left-lateralized Stroop system, but that it reflects a developmentally early form of prefrontal organization.

## Methods

### Participants and procedures

Ninety-four children aged 35–79 months were analyzed. Participants were recruited from kindergartens and nursery schools in Osaka, Japan. Informed consent was obtained from caregivers and preschools before the children were involved in the study, according to the principles of the Declaration of Helsinki. This study was approved by the ethics committee of the Unit for Advanced Studies of the Human Mind at Kyoto University. An additional five children were excluded: four were unable to complete the task at the beginning of the experiment, and one refused to wear the fNIRS band. No diagnosis of a developmental disorder had been reported for any of the children. Additional participant information is provided in Supplementary Note 1.

### Sample size

Neuroimaging and developmental studies, including fNIRS studies using the Stroop task, have frequently been limited by small samples and insufficient statistical power^53,58^. We therefore designed the study with a large, reliable sample as a primary objective. An a priori power analysis conducted in G^*^Power (v. 3.1) indicated that a sample of *n* = 84 was required to detect a moderate correlation (*r* = 0.30) with 80% power at α = 0.05 (two-tailed). The analyzed sample of *n* = 94 exceeded this requirement. Although this study was not formally pre-registered, the sample size was determined a priori on the basis of this correlation-based power analysis, and the linear mixed-effects model approach was planned and adopted, in line with prior studies^31^, as a method suited to comprehensively capturing region-specific brain activity.

### Stroop-like task

Following the method described by Moriguchi^40^, children performed a Stroop-like conflict task. Specifically, this task was a modified pointing version of the Black/White task^55,59^, while retaining the basic Stroop-like conflict structure shared with the Grass/Snow task^6^. Because the classic color-word Stroop task places demands on reading and automatic word recognition, which vary with reading skill during development^35^, we used an age-appropriate color-pointing Stroop-like variant. In this task, children listened to spoken color labels and responded by pointing, rather than reading words or producing verbal color names. Three color pairs (red/blue, black/white, and yellow/green) were printed on A3-sized laminated cards (Fig. 1). Each card displayed two colored rectangles side by side against a gray background. Each color pair was administered under two conditions (congruent and incongruent), yielding six sessions in total (3 pairs × 2 conditions). In the congruent condition, the experimenter named a color aloud and the child pointed to the card of the matching color (e.g., when the experimenter said "red," the child pointed to the red card) (37.2 ± 10.6 trials in total, mean ± *SD*). In the incongruent condition, the child was required to point to the card of the opposite color (e.g., when the experimenter said "red," the child pointed to the blue card) (32.6 ± 10.7 trials in total, mean ± *SD*). This reverse-contingency rule required the child to suppress the prepotent response to the named color and select a rule-based opposite-color response, creating a Stroop-like conflict structure rather than a direct instantiation of the classic color–word Stroop task. Within each session, trials of the two colors were presented in a pseudorandom order. The condition order was fixed across all children, with the congruent condition always preceding the incongruent condition. For the youngest children in particular, it was essential that they first acquire the basic task operation (pointing to a card in response to the experimenter’s color instruction) before being introduced to the reverse rule. Each session consisted of a 15-s rest phase followed by a 25-s task phase. The primary dependent variable was response accuracy (number of correct responses / total number of completed trials) in each condition, following previous studies^40^.

### fNIRS recording

Prefrontal hemodynamic responses were recorded using a 16-channel continuous-wave fNIRS system (OEG-16; Spectratech Inc., Tokyo, Japan) emitting near-infrared light at two wavelengths (770 nm and 840 nm). The probe configuration followed the protocol established and validated in our previous developmental fNIRS study of executive function in preschool children^31–33,40^. The probe array comprised six light-emitting and six detector optodes, forming 16 channels at a source-detector separation of 30 mm. The probe was positioned with the lower row centered on Fpz, following the same configuration as in our previous study^31^. Channel locations and anatomical coverage were established in prior work from our group using spatial registration to MNI standard space and probabilistic anatomical labeling^31^; upper-row channels covered primarily the DLPFC (Brodmann areas 9/46) and lower-row channels covered primarily the RLPFC (lateral Brodmann area 10, frontopolar region) (Supplementary Note 2 and Supplementary Tables 1 and 2). At acquisition, valid fNIRS values were obtained only from channels that passed the pre-measurement calibration implemented in the OEG-16 acquisition software. Raw optical data were converted to hemoglobin concentration changes using the modified Beer-Lambert law. Relative changes in oxy-Hb were calculated in units of mM·mm without assuming specific differential pathlength factors^60^.

### fNIRS preprocessing

Data were sampled every 0.655 s and preprocessed and analyzed using the OEG-16 proprietary software. The fNIRS waveforms were reviewed alongside the corresponding video footage of each child’s behavior by trained technicians and researchers to identify excessive motion-related noise, according to previously described procedures^31^. Exclusions were minimal leaving 99.3% of the measured trials and 99.6% of the measured channels for analysis. Data were smoothed with a seven-point moving average filter and subsequently detrended using linear fitting to reduce slow drift. Hemodynamic modality separation^57^ was then applied to each channel to decompose the signal into a functional component and a systemic component. This approach exploits the principle that task-evoked cortical activation generates a negative relationship between oxy-Hb and deoxy-Hb, whereas systemic fluctuations and motion artifacts produce positively correlated changes in both signals. Consistent with prior applications in young children from our group^31,33,40^, this procedure yielded reliable estimates of task-evoked functional activity. Subsequent analyses used the functional oxy-Hb component, which provides a higher signal-to-noise ratio and more accurately reflects cortical hemodynamic responses. ROI signals were computed by averaging across channels covering each region: rDLPFC (ch 2, 4, 5), lDLPFC (ch 11, 13,14), rRLPFC (ch 3, 4, 6), and lRLPFC (ch 12, 13, 15). Channels spanning two ROIs (ch 4, 13) were included in each ROI with a weighting of 0.5. Left DLPFC channels that did not pass the pre-measurement calibration were unavailable for five participants (*N* = 89; Supplementary Note 1). Task-evoked activity was quantified as the mean oxy-Hb change during the task window (4.6 s after onset to 15 s after task offset) relative to a pre-task baseline (3.3 s to 0 s before onset)^31^. A sequential Grubbs test (α = 0.0001) was applied to mean oxy-Hb values within each ROI; no outliers were identified.

### Analysis plan

Statistical analyses were conducted in R (version 4.3.2). The significance threshold was set at α = 0.05. Accuracy was compared between conditions using Wilcoxon signed-rank tests, and Pearson correlations with age were computed for accuracy in each condition and for the interference score (incongruent minus congruent). Oxy-Hb responses were analyzed using linear mixed-effects models (lme4, lmerTest,emmeans), with fixed effects of condition (congruent, incongruent), hemisphere (left, right), region (DLPFC, RLPFC), and random intercepts for participants. Main effects and interactions were assessed using Type III ANOVA with Satterthwaite approximation, followed by post-hoc pairwise comparisons. To test for developmental modulation, age was added to the model as a main effect and as an interaction with condition; sex was tested as an additional covariate. To examine whether the Stroop-like effect was present across the full age range, condition effects were estimated at representative ages (36, 48, 60, 72 months) using emmeans. To examine brain–behavior relationships, Pearson correlations and age-partialled partial correlations were computed between ROI oxy-Hb responses and response accuracy, for congruent, incongruent, and interference (incongruent minus congruent) measures, separately. Where multiple comparisons were performed across four ROIs, false discovery rate (FDR) correction was applied within each set of comparisons, and both uncorrected and FDR-corrected P values are reported. As an exploratory analysis, lateralization indices were computed as right minus left oxy-Hb for each ROI following the same hemispheric-difference approach used to index frontal laterality in fNIRS Stroop studies^21,22^.

## Acknowledgements

The authors thank Dr. Watanabe (Kyoto University) for valuable discussions and the technical and administrative assistants in the laboratory for their dedicated support throughout the study. The authors also express their gratitude to Ms. Noguchi (ELCS–English Language Consultation Services, Japan) for assistance with the manuscript. We are deeply grateful to the staff of the participating kindergartens and nursery schools, as well as to all child participants and their guardians, for their generous cooperation. The illustration of the girl in Fig. 1a was drawn by Atsuko K.

## Source of funding

R.K. was supported by JSPS KAKENHI (Grant Numbers: 23H04830, 24K20598, and 23KJ1169). Y.M. was supported by JSPS KAKENHI (Grant Numbers: 24K00486, 23H04832). N.Y., and R.K. also acknowledge the financial support from a crowdfunding project through the academic crowdfunding platform "academist" (Project No. 396).

## Author contributions

**Ryuta Kuwamizu:** Conceptualization, Methodology, Performing the experiments, Data analysis, Visualization, Writing—Original draft preparation, Project administration, Funding acquisition. **Nozomi Yamamoto**: Writing—Reviewing and Editing, Participants recruitment, Funding acquisition. **Kota Otani:** Data analysis support, Writing—Reviewing and Editing. **Yusuke Moriguchi**: Conceptualization, Supervision, Writing—Reviewing and Editing, Funding acquisition.

## Competing interests

The authors do not declare any competing interests.

## Data availability

The repository will be made publicly available upon publication.

## Notes

### Competing Interest Statement

The authors have declared no competing interest.

