## Supplementary Information for "Right-lateralized prefrontal signature with successful Stroop-like interference control in early childhood"

**Supplementary Note 1. Additional participant information**

Ninety-four children (48 boys, 46 girls) aged 35–79 months (mean = 58.7, SD = 11.1) contributed data. Children completed 37.2 ± 10.6 congruent and 32.6 ± 10.7 incongruent trials (mean ± SD). Accuracy was near ceiling on congruent trials (0.954 ± 0.053; median 0.970) and lower and more variable on incongruent trials (0.750 ± 0.260; median 0.845; range 0.00–1.00), giving a mean interference score of −0.203 ± 0.241. Participant-level values for all variables are provided in the source data file.

Because melanin absorbs near-infrared light, hair colour can attenuate the fNIRS signal[1], and it was therefore recorded on a four-point scale: black, 36 (38.3%); dark brown, 47 (50.0%); brown, 8 (8.5%); light, 3 (3.2%). The sample was predominantly dark-haired (black or dark brown, 88.3%). Controlling for hair colour did not alter the associations reported in the main text: incongruent oxy-Hb remained correlated with incongruent accuracy in the right DLPFC ($r = 0.355$, $P < 0.001$) and the right RLPFC ($r = 0.349$, $P < 0.001$), as did the RLPFC lateralization index with incongruent accuracy ($r = 0.302$, $P = 0.003$) and with the interference score ($r = 0.291$, $P = 0.005$). These are partial correlations controlling for hair colour, which was entered as an ordinal covariate (0 = black to 3 = light); P values are uncorrected.

Hand use during the pointing response was recorded on a five-point scale: right, 66 (70.2%); mostly right, 5 (5.3%); both hands equally, 11 (11.7%); mostly left, 3 (3.2%); left, 9 (9.6%). Seventy-one children (75.5%) used the right hand exclusively or predominantly. Controlling for hand use likewise did not alter these associations: the right DLPFC ($r = 0.364$, $P < 0.001$) and the right RLPFC ($r = 0.345$, $P < 0.001$) remained correlated with incongruent accuracy, as did the RLPFC lateralization index with incongruent accuracy ($r = 0.308$, $P = 0.003$) and with the interference score ($r = 0.299$, $P = 0.004$). Hand use was entered as an ordinal covariate (0 = left to 4 = right); P values are uncorrected.

Valid fNIRS values were obtained only from channels that passed the OEG-16 pre-measurement calibration; channels that did not pass were not measured, so no signal was acquired for them. Left DLPFC values and the corresponding DLPFC lateralization indices were consequently unavailable for five participants (N = 89) and were analyzed as available cases without imputation. Oxy-Hb values are expressed in mM·mm. DLPFC, dorsolateral prefrontal cortex; RLPFC, rostrolateral prefrontal cortex; oxy-Hb, oxygenated hemoglobin.

Participants also completed another cognitive task during the same study visit; data from that task for a subset of the present sample have been reported elsewhere[2].

**Supplementary Note 2. fNIRS channel localization and ROI assignment**

The fNIRS probe montage and region-of-interest (ROI) definitions used in the present study were identical to those used in our previous developmental fNIRS study with the same OEG-16 configuration in preschool children. Accordingly, channel localization and probabilistic anatomical labeling followed the previously established registration procedure reported by Kuwamizu et al.[2]. Briefly, spatial registration was conducted in an independent sample of six preschool children (2 female; mean age = 55 months, SD = 11.9, range = 42 –72 months) recruited from the same age cohort. Head surfaces were captured using a Revopoint MIRACO Pro 3D scanner, and optode/channel locations were reconstructed with OEG-3DXYZ-OBJ (Spectratech, Japan). Channel positions were registered to MNI standard space using NIRS-SPM[3], and probabilistic anatomical labeling was obtained using NFRI functions.[4] Brodmann area information was summarized with MRIcro[5]. Because the present study used the identical montage and ROI assignment, the previously established MNI coordinates and ROI-level anatomical labels are reproduced here.

**Supplementary Table 1.** MNI coordinates for fNIRS channels. Spatial registration was based on an independent sample of six preschool children. The table reports estimated MNI coordinates (x, y, z; mean ± SD across participants) for channels used in the ROI definitions. The localization information follows Kuwamizu et al.[2].

| Channel | MNI coordinates (mm) | | |
|---|---|---|---|
| | x | y | z |
| Ch2 | 45.5 ± 3.6 | 45.2 ± 3.4 | 28.7 ± 4.5 |
| Ch3 | 45.0 ± 3.8 | 57.5 ± 4.8 | 5.3 ± 5.9 |
| Ch4 | 37.0 ± 5.0 | 59.8 ± 3.3 | 19.3 ± 5.2 |
| Ch5 | 25.0 ± 4.8 | 59.5 ± 3.3 | 32.2 ± 5.3 |
| Ch6 | 25.2 ± 4.5 | 69.8 ± 1.8 | 10.3 ± 6.4 |
| Ch11 | -23.0 ± 4.0 | 57.8 ± 4.8 | 33.5 ± 4.3 |
| Ch12 | -25.5 ± 4.6 | 68.0 ± 3.3 | 10.0 ± 5.6 |
| Ch13 | -36.8 ± 5.0 | 57.2 ± 5.8 | 20.3 ± 4.6 |
| Ch14 | -44.5 ± 4.1 | 40.7 ± 7.8 | 30.7 ± 4.0 |
| Ch15 | -46.5 ± 4.2 | 51.5 ± 6.3 | 4.7 ± 5.5 |

**Supplementary Table 2.** ROI probabilistic anatomical labels. Probabilistic anatomical labeling indicates the relative overlap probability of each ROI with anatomical regions (N = 6). ROI-level labeling distributions were computed using the same channel weights as used for ROI signal computation. Boundary channels are shown in parentheses and were weighted as 0.5. Abbreviations: DLPFC, dorsolateral prefrontal cortex; RLPFC, rostrolateral prefrontal cortex; FPA, frontopolar area; OFA, orbitofrontal area; IFG, inferior frontal gyrus; r, right; l, left. ROI definitions follow Kuwamizu et al.[2].

| ROI | Channel | Region (Brodmann area) [%] | Top label |
|---|---|---|---|
| rDLPFC | Ch2, (Ch4), Ch5 | DLPFC (9/46) [58.7%], Pars triangularis (45) [21.5%], FPA (10) [19.8%] | DLPFC (9/46) |
| lDLPFC | Ch11, (Ch13), Ch14 | DLPFC (9/46) [58.9%], Pars triangularis (45) [24.4%], FPA (10) [14.6%], Pars opercularis (44) [2.1%] | DLPFC (9/46) |
| rRLPFC | Ch3, (Ch4), Ch6 | FPA (10) [56.2%], DLPFC (9/46) [37.2%], OFA (11) [5.8%], Pars triangularis (45) [0.8%], IFG (47) [0.1%] | FPA (10) |
| lRLPFC | Ch12, (Ch13), Ch15 | FPA (10) [43.4%], DLPFC (9/46) [40.9%], Pars triangularis (45) [10.5%], OFA (11) [5.1%] | FPA (10) |
